# Multichannel bioelectronic sensing using engineered *Escherichia coli*

**DOI:** 10.1101/2023.09.30.560307

**Authors:** Xu Zhang, Caroline Ajo-Franklin

## Abstract

By engineering extracellular electron transfer (EET) to be dependent on an analyte, researchers have developed whole cell bioelectronic sensors that sense hazards to human and environmental health^1^. However, these sensors regulate a single electron transfer pathway as an electrochemical channel, limiting the sensing information to a single analyte. To increase information content, we developed a multichannel bioelectronic sensor through which different chemicals regulate distinct extracellular electron transfer pathways within a single *Escherichia coli* cell. One channel utilizes the flavin synthesis pathway from *Bacillus subtilis*^2^ and the other a set of cytochromes constructing the Mtr pathway from *Shewanella oneidensis*^3^. We demonstrate an arsenite responsive promoter can control the Mtr pathway through activation of cytochrome CymA expression and a cadmium responsive promoter can control the flavin synthesis pathway^4,5^. The redox potential of flavin-mediated EET is different from that of CymA-mediated one^6^. This allowed for development of a redox-potential-dependent algorithm that distinguishes variable input signals of each analyte mediated by two EET pathways in vivo. This approach enables a 2-bit binary signal readout for real-time tracking throughout the entire sensing duration. Our multichannel bioelectronic sensor was able to accurately sense and distinguish different heavy metals in Brays Bayou water samples with a response time comparable to that in clean water. This multichannel bioelectronic sensors allow for simultaneous detection of different chemicals, significantly expanding information transmission and helping to safeguard human and environmental health.

## INTRODUCTION

Environmental water contamination, stemming from both natural and synthetic chemicals, poses a significant global challenge. While sophisticated analytical techniques such as ion chromatography^7^ or atomic absorption spectroscopy^8^ can provide accurate results even in complex samples, they still rely on manual collection, controlled storage, and long-distance transportation of water samples back to a laboratory. These preparation and sampling procedures are expensive and time-consuming^9^, thereby limiting the frequency of analysis. This lack of time resolution is prone to missing short-lived events, such as periodic releases of toxins or pollutants, which can have significant consequences to environmental and human health.

In contrast to lab-based analytical techniques, in situ chemical sensors allow for more frequent monitoring, even in remote locations^10^. However, this convenience comes at the expense of selectivity in complex conditions^11,12^. To enhance the selectivity of in situ chemical sensors, preliminary sample separation steps, such as chromatography and electrophoresis are incorporated^13^. However, these preparation steps require substantial power and bulkier instrumentation.

Biosensors, which rely on biomolecular recognition, are an alternative to chemical sensors. They can be highly selective, portable and have minimal power requirements^14^. Enzymatic biosensors stand out due to their remarkable selectivity for a specific substrate and rapid response time^15^. However, enzymatic biosensors face certain drawbacks, such as the high cost associated with producing purified enzymes and short operational lifespan^16–19^. In contrast, whole cell biosensors exploit the self-maintenance and self-replication capabilities of living microorganisms resulting in lower cost and longer lifespan times^20–23^.

Bioelectronic sensors, a type of whole cell biosensor, employ electroactive microorganisms to transduce chemical signals into readable electrical signals^1,24^. The transduction process that enables microbes to deliver electrons from the intracellular donor to extracellular acceptor is called extracellular electron transfer (EET). Synthetic biology allows us to construct sensing components to control the EET process, enabling bioelectronic sensing of desired targets^25^. Recently developed bioelectronic sensors are capable of sensing endocrine disruptors with response times on the minute scale^1^, but since they employ only one EET pathway within the whole cell sensor, they are limited to detection of a single analyte. Monitoring environmental water quality requires measuring multiple parameters simultaneously for a complete understanding of contamination and thus higher spatio-temporal resolution of must be engineered into bioelectronic sensors.

To increase the amount of sensing information, we developed a bioelectronic sensor with two EET pathways in a single strain of *Escherichia coli*. The system allows for sensing of two common environmental pollutants, arsenic (As) and cadmium (Cd) simultaneously. By employing two distinct heavy metal responsive promoters, the two EET pathways are independently activated in response to the presence of As or Cd, leading to increases in current values. To distinguish between the 4 possible states based on the presence or absence of each analyte, we developed a redox-potential-dependent algorithm that effectively encodes the signals into 2-bit binary outputs corresponding to the input signals. Utilizing this algorithm, we can accurately identify the analytes within both purified water or environmental water samples, as well as conduct both single-measurement or continuous monitoring.

## RESULTS

To create a multichannel bioelectronic sensor, we were inspired by how light signals are sensed and processed by the color vision system^26,27^ (Fig 1 top). Visual light is composed of a spectrum of colors, each associated with a specific wavelength. These wavelengths are first sensed by three different receptors and converted into electrical signals, which then travel through separate channels to finally reach a shared bipolar ganglion cell. This cell performs an algorithmic computation on these signals through the opponent process and outputs the results to the visual cortex, which comprehends the mix of wavelengths as color^28^.

**Fig 1.**
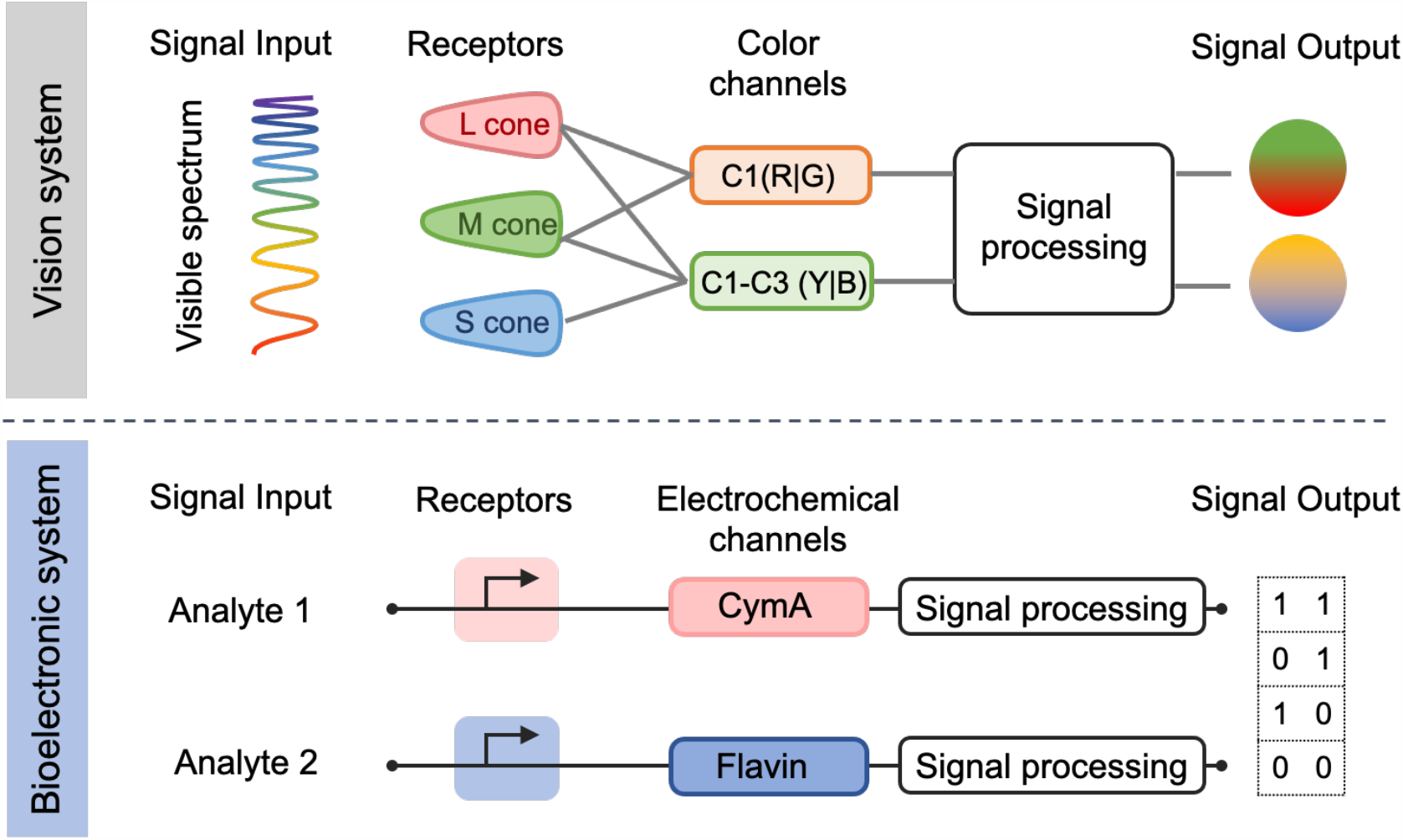
The process of a multichannel bioelectronic sensor system can be analogized to that of the color vision system. In both systems, specialized channels are triggered by input signals, which are then transduced and processed into either the final optical signal or digitized signals.

Similar to the color vision system, we engineered a single strain of *E. coli* with two EET pathways controlled by specific analytes and transformed the biological responses into electrical signal output. To achieve this, we have constructed transcriptional regulation systems capable of sensing target analytes and activating expression of distinct EET pathways. The EET pathways then conduct electrons from the cytoplasm to electrodes to generate electrical signals at different redox potentials, referred to as electrochemical channels. These signals from each electrochemical channel are recorded by a potentiostat-controlled electrode, processed by a voltammetric algorithm, and encoded into 2-bit binary signal output. Each digit corresponds to signal or lack thereof from individual channels and thus, the various combinations of two analytes can be represented by four different 2-bit binary signals.

### Design strategy for EET pathways

As the essential signal transduction channels, the two EET pathways need to be engineerable, produce measurable current, and have different redox potentials. The CymA limited MtrCAB pathway (CymA pathway for short) from *Shewanella oneidensis* and flavin synthesis pathway from *Bacillus subtilis* were selected because both have been successfully engineered into *E. coli*^2,6,29^, and the activation of each individual EET pathway delivered measurably higher current signals than the basal signal associated with the wild type *E. coli*. Additionally, the redox potentials (E_1/2_) associated with the CymA and Flavin pathways are differentiable (0 mV vs -400 mV vs. Ag/AgCl)^6,29,30^. Thus, these pathways formed the basis for our sensing channels.

### Constructing Two Electron Transfer Pathways in *E. coli* as Sensing Channels

To enable multiplexed sensing, we employed two different types of signal receptors coupled to the EET pathways. We first selected commonly used inducible promoters to individually sense and activate the two EET pathways and refer to this strain as ***inducer-E. coli*** (Fig 2A and 2D, middle). Specifically, an IPTG-activated promoter regulates the expression of the *cymA* gene while a constitutive promoter controls the *mtrCAB* genes from *Shewanella oneidensis*^29^. An aTc-activated promoter controls the *ribADEHC* genes responsible for flavin synthesis from *Bacillus subtilis* as the flavin synthesis pathway^25^. To demonstrate applicability in environmental monitoring^4,5^, we incorporated heavy metal responsive promoters to detect multiple metals simultaneously and refer to this strains as ***heavy metal-E. coli*** (Fig 2A and 2D, right). In the heavy metal-*E. coli*, a modified arsenite-regulated promoter was used to activate CymA expression in response to the presence of As^5^, while flavin synthesis is regulated by a cadmium-inducible promoter to sense Cd presence^4^. The subsequent electrical signals delivered via the different EET pathways are thus referred to as the **CymA-Channel** and the **Flavin-Channel**.

**Fig 2.**
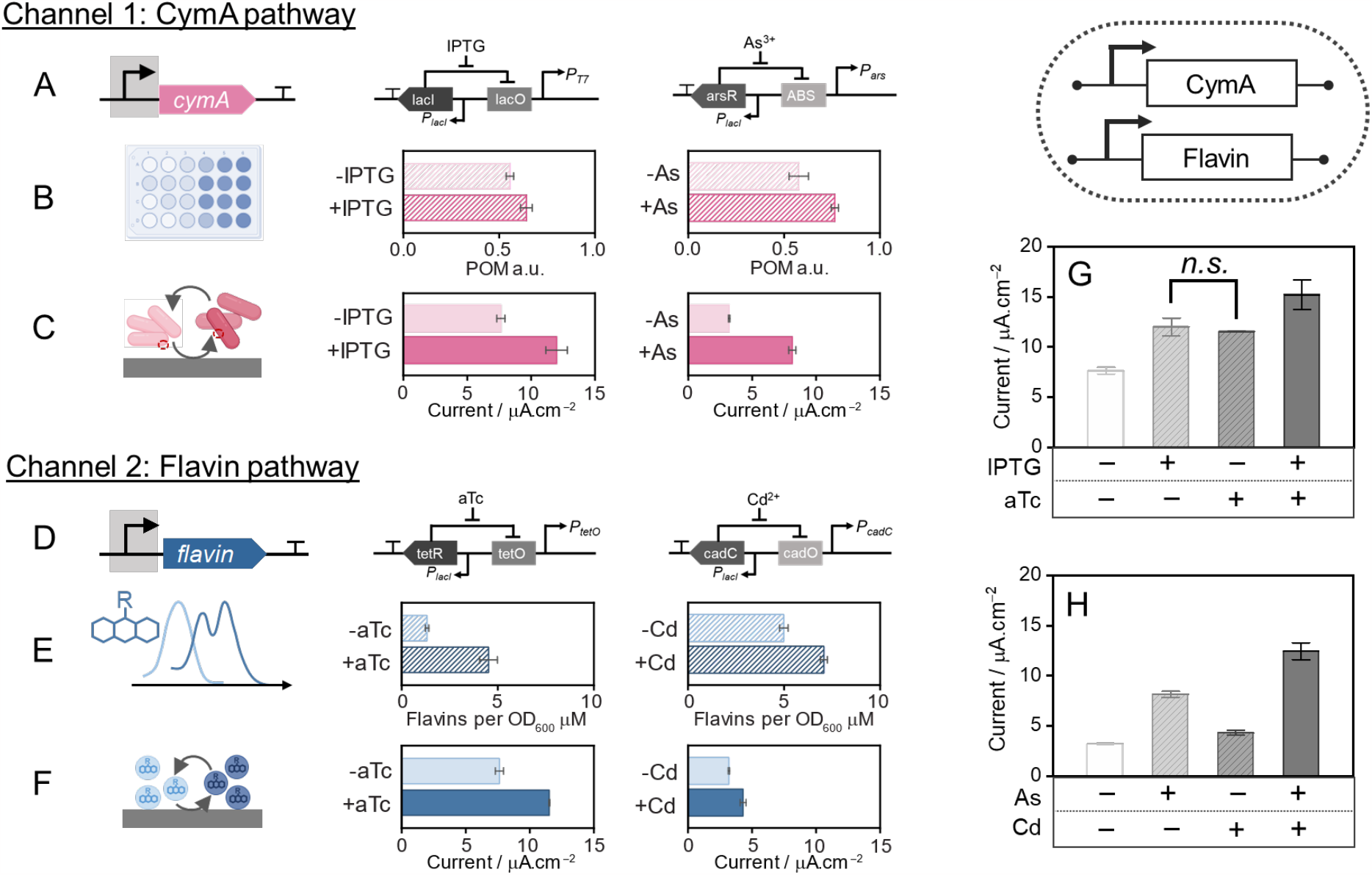
An *E. coli* sensor with multiple electron transfer pathways as individual and hybrid channels readout. (**<u>A-C</u>**) The CymA-limited pathway from *Shewanella oneidensis* was genetically engineered into *E. coli* with inducible promoters to modulate extracellular electron transfer (EET) activation, henceforth referred to as CymA pathway. (**A**) A schematic diagram illustrating the genetic modules containing inducible promoters to activate *CymA* expression to control EET activity (left). The genetic modules utilized either a IPTG-inducible T7 promoter (middle) or an arsenic-responsive promoter (right). (**B**) CymA-limited direct electron transfer was evaluated using a colorimetric molybdenum-based nanoparticles assay, where the increase of blueness corresponded to higher EET activities (schematic, left). The colorimetric POM intensity in the presence and absence of induction by 100 μM IPTG (middle) or 2.5 μM As (right), showing a greater EET response with IPTG and arsenic induction. (**C**) Electrical signals were measured by culturing cells with continuously polarized electrodes at +0.20 V vs. Ag/AgCl (schematic, left). Steady-state current in the presence and absence of induction by IPTG (middle) or As (right), showing a greater bioelectrochemical response with IPTG and As induction. (**D-F**) The flavin synthesis pathway from *Bacillus subtilis* was engineered using inducible promoters to modulate mediator-based EET activity. (**D**) A schematic diagram illustrating the genetic modules containing inducible promoters to activate flavin synthesis to control mediated EET activity (left). The genetic modules utilized either aTc-inducible tet promoter (middle) or a Cd-responsive promoter (right). (**E**) Flavin-mediated electron transfer was investigated using the intrinsic fluorescence of flavins. The fluorescence intensities per OD were higher in the presence of 4 μM aTc (middle) or 800 nM Cd (right) compared to conditions without aTc or Cd. (**F**) Electrical signals were measured by culturing cells with continuously polarized electrodes at +0.2 V vs Ag/AgCl (schematic, left). Steady-state current in the presence and absence of induction by aTc (middle) or Cd (right), showing a greater bioelectrochemical response with aTc and Cd induction. (**G-H**) To analyze our ability to distinguish between the bioelectrochemical response to no analytes, one analyte, or both analyte, both the CymA channel and Flavin channel were measured in tandem. (**G**) Steady-state current response from E. coli in the presence of different combinations of aTc and IPTG (**H**) Steady-state current response from E. coli in the presence of different combinations of As and Cd In both G-H, it is challenging to discriminate between all four conditions based on the steady-state current level. For all POM assay and flavins characterizations, data represent the mean values, with error bars representing one standard deviation (n = 4 biologically independent samples). For the electrical current measurements in bioelectrochemical systems as G and H, the data represents the mean values, with error bars representing one standard deviation (n=3 biological independent samples).

To optimize the function of the individual CymA-Channel and Flavin-Channel, we assayed the expression level of the CymA pathway and Flavin synthesis pathways respectively as a factor of induction level. To assess the expression of the CymA pathway, we incubated the *inducer-E. coli* or the *heavy metal-E. coli* with cell-impermeable molybdenum-based nanoparticles, a more sensitive particle for reporting on EET than commonly used WO_3_^31^. With this assay, the nanoparticles undergo an electrochromic shift from white to blue based on the level of reduction by CymA-channel. The cells were then induced at different levels with IPTG (<1000 μM) or As (<100 μM) and the color change of the nanoparticles recorded (Fig S2). The EET performance of the CymA-Channel is substantially enhanced when induced at 100 μM IPTG or 2.5 μM As compared to non-induced cells (Fig 2B). Thus, for all subsequent studies, we induced cells with 100 μM IPTG or 2.5 μM As to maximize EET activity while minimizing toxicity to cells (Fig S2).

The flavin synthesis pathway was characterized based on the intrinsic fluorescence properties of flavins. As redox mediators, flavins can deliver the electrons from the interior of the cell to the electrode and generate electrical signals. The production of extracellular flavins peaked with induction at 4 μM aTc and was substantially improved after induction with 800 nM Cd (Fig 2E). Thus, for all subsequence experiments on the flavin synthesis pathway, we induced cells with 4 μM aTc or 800 nM Cd.

After using assays to assess the expression level of the CymA and Flavin synthesis pathways, we inoculated each engineered strain into bioelectrochemical systems (BES) to assess the *in situ* electrical signal delivery^1^. We directly added both cells and analytes into the bioreactors. When the cells were exposed to one of the analytes, the expression of the corresponding EET pathway was activated and led to a current increase (Fig 2C, 2F), demonstrating that each individual EET pathway is functional as a single electrochemical channel. Interestingly, the inducer-*E. coli* synthesizes less flavins compared to heavy metal-*E. coli*, while achieving a higher current output (Fig 2C, 2F). This might be because of the higher basal current output from inducer-*E. coli*, possibly due to the inefficient transcriptional control of the IPTG promoter. The promoter ‘leakage’ led to higher background expression of the CymA pathway, resulting in higher current level even without adding IPTG^32^. As a consequence, when the Flavin-channel was activated by the presence of aTc, the inducer-*E. coli* produced a greater current output due to the addition of this background current.

To determine whether our bioelectronic sensor can be used as a two-channel biosensor, we induced the strains with different combinations of analytes: none of analytes, either of analytes, or both of analytes. While some combinations were easily differentiated, in the inducer-*E. coli*, the current level activated by IPTG only is 11.0 μA.cm^−2^, which is close to the 11.5 μA.cm^−2^ level activated by aTc only (Fig 2G, current evolution curves shown in Fig S3). Also, when we evaluate the electrical signal output of the heavy metal-*E. coli*, the current level activated by Cd only is quite close to the basal current without any heavy metals present (Fig 2G). These similar current levels collected at a single potential across various analyte conditions (e.g., IPTG-only vs. aTc-only; uninduced control vs. Cd-only shown in Fig 2H and 2G) pose a challenge to accurately identifying the different combinations of analytes.

### Developing a Redox-Potential-Dependent Algorithm for Signal Processing

To overcome the challenge of signal differentiation, we developed an algorithm that relies on the distinct redox potentials (E_1/2_) associated with the two EET channels. Since the redox potential of the CymA channel is more positive than that of the Flavin channel, we evaluated the oxidation process of the CymA channel and the reduction process of the Flavin channel.

We applied two different potentials sequentially to the electrode over a 30-minute time decay (shown in Fig 3A and 3B), a technique referred to as double potential step chronoamperometry (DPSC). The two potentials were employed to create two distinct conditions: one for basal level and another for either oxidation dominance by the CymA channel or reduction dominance by the Flavin channel. The current ratios between the two potentials were obtained and then normalized with respect to the basal condition in the absence of any analytes (Fig 3C, 3D). These ratios are associated with the expression levels of each EET pathway and the relative current ratios as the function of time decay are depicted in a heatmap. The CymA channel is depicted in red, while the Flavin channel is presented in blue. The darkness of the colors corresponds to the values of the normalized current ratios.

**Fig 3.**
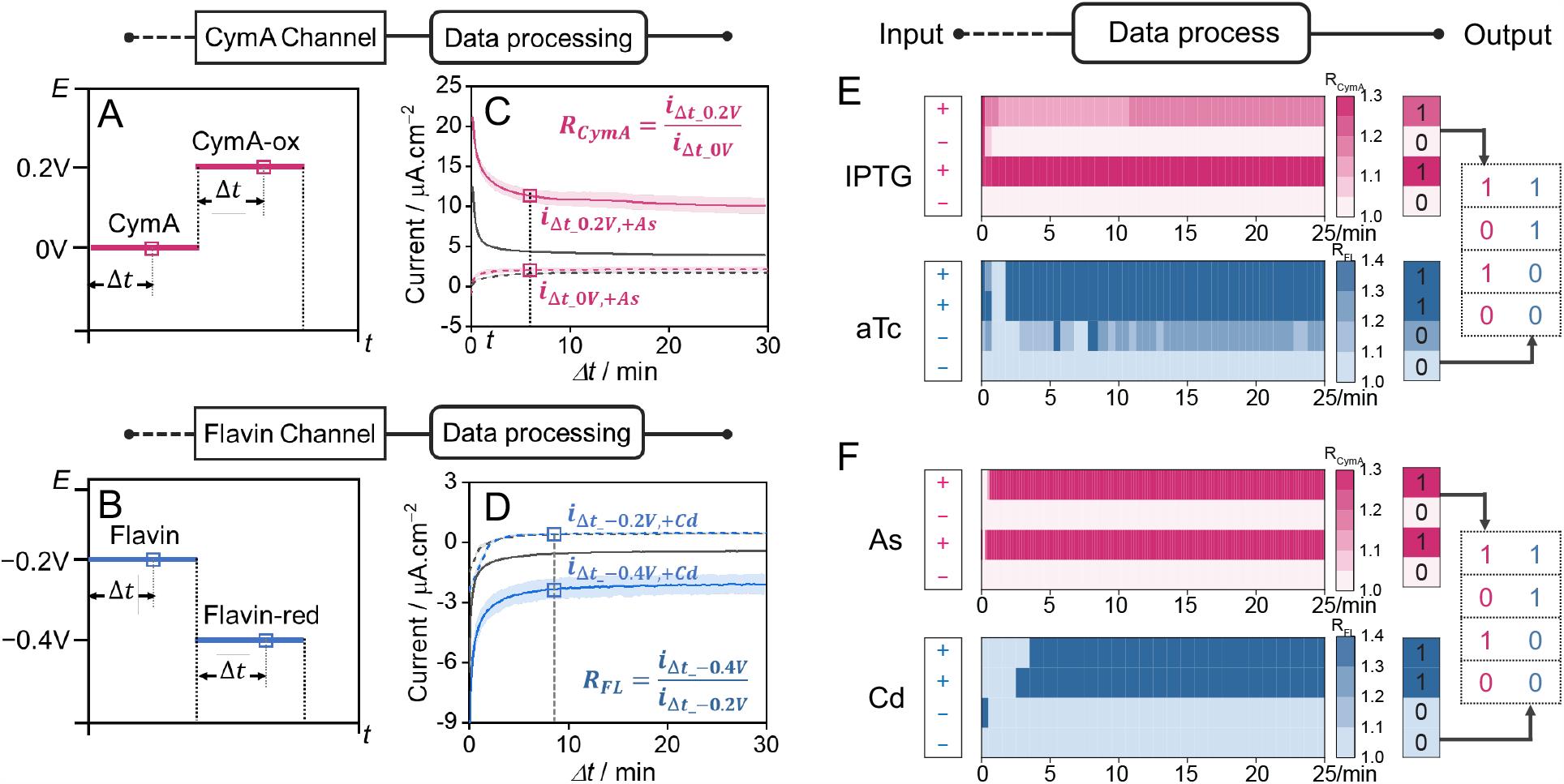
The two electrochemical channels were differentiated via a voltammetric algorithm based on the distinct redox potentials of the CymA and Flavin synthesis pathways. (**A-B**) Schematic diagram depicting the waveform for the double potential step experiment to assess the electrical flux mediated by (**A**) CymA Channel or (**B**) Flavin Channel. The first potential applied is more positive than the midpoint potential so that the redox center is fully oxidized, while the second potential step applied is lower than the midpoint potential, allowing re-reduction of the redox center. For each step, the current is measured from the start of the potential step by a time delay (Δt). (**C**) Current as a function of delay time for the CymA channel with the first potential set to 0.00 V (dashed lines) and the second potential set to +0.20 V (solid lines). Current was measured for heavy metal-E. coli strain in the absence (gray lines) or presence of As (red lines). (**D**) Current as a function of delay time for the RF channel with potential steps of −0.20 V (dashed lines) and −0.40 V (solid lines). Current was measured for what strain in the absence (gray lines) or presence of Cd (blue lines). The currents from the first and second potential step at a fixed time delay were used to generate the current ratio, R_CymA_ or R_RF_, as shown in the equations. (**E-F**) Heatmaps showing R_CymA_ (red) or R_RF_ (blue) as a function of time delay. (**E**) Heatmap for inducer-E. coli strain in the presence and absence of IPTG and aTc. (**F**) Heatmap for heavy metal-E. coli in the presence and absence of As and Cd. The current ratio was then normalized with respect to the absence of both analytes in order to plot the heatmaps. Heatmap color intensity varies by ratio values to show analyte presence (1, dark red/blue) or absence (0, light red/blue). For each channel, the digitized signals were signed based on the threshold of current ratios 1 (CymA > 1.15, Flavin > 1.2) or 0.

We observed a strong correlation between the presence or absence of analytes and the color patterns displayed on the heatmaps (Fig 3E, 3F). For both CymA and Flavin channels, the presence of analytes (marked as ‘+’) indicated by higher current ratios corresponds to darker shades in the heatmaps and is assigned a ‘1’ when the ratio is higher than the thresholds (CymA > 1.15, Flavin > 1.2). Conversely, in absence of analytes (marked as ‘–’) lower current ratios were calculated and thus lighter colors were observed on the heatmaps and assigned a ‘0’ (Fig 3E, 3F). Hence, a 2-bit binary signal configuration was implemented wherein four states are possible. The first digit was dedicated to indicating the signals associated with the CymA channel, specifically responding to the presence or absence of IPTG or As. The second digit encoded the signals attributed to the Flavin channel in response to the presence or absence of aTc or Cd. This configuration allows for the encoding of information related to the activation of the two EET channels in a concise and distinct manner.

Moreover, this configuration can be optimized into a shorter time frame for analytes detection. For the CymA channel, it takes 10 minutes for the algorithm to demonstrate the presence of IPTG while it takes 1 minute for As recognition. For the Flavin channel, it takes 2 minutes for the algorithm to determine the presence of aTc while it takes 3 minutes for Cd recognition. (Fig 3E and 3F). This fast detection allows a shorter polarization time of each potential instead of 30 min. In light of this, for the rest double potential step chronoamperometry, we have decreased the time duration for each potential to 15 min.

### Continuous Surveillance of Diverse Analyte Combinations in Environmental Water Samples

After demonstrating the double potential analysis works to differentiate the signals for a single measurement, we applied it for multiple measurements over 40 hours, enhancing the temporal resolution of bioelectronic sensing measurements (Fig 4, current evolution with sampling time points shown in Fig S5). In purified water samples, both the inducer-*E. coli* and heavy metal-*E. coli* bioelectronic sensors were able to continuously distinguish the various combinations of analytes over approximately 40 hours of testing (Fig 4B, 4C). While utilizing the inducer-*E. coli* strain, we were only able to detect IPTG presence after 18 hours, but we were able to identify all 4 different combinations of analytes at the 30-hour mark (Fig 4B). Similarly, when employing the heavy metal-*E. coli* strain, we detected the presence of As or Cd alone after 18 hours and identified the other conditions at the 30-hour mark (Fig 4C).

**Fig 4.**
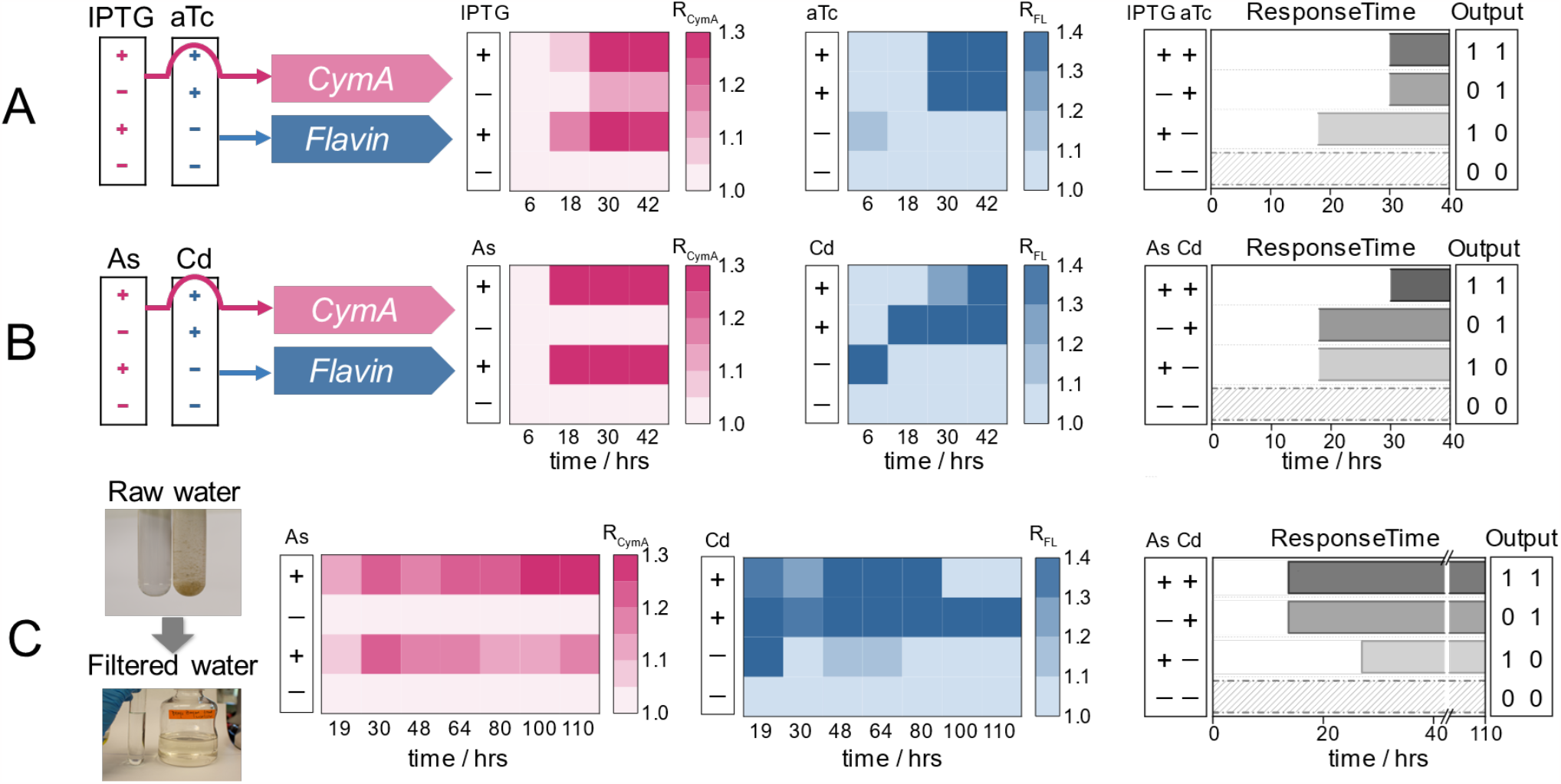
Dynamic monitoring of algorithmically processed signals in both laboratory conditions and environmental water samples collected from Brays Bayou, Houston. (**A-B**) Schematic depicting the multichannel bioelectronic sensing process in clean water. Inducible modules were utilized to activate cymA or flavin synthesis gene expression through different regulated factors: IPTG or aTc activation system (A) and As or Cd activation system (B). (**C**) Schematic displaying the multichannel bioelectronic sensing process to detect heavy metals As or Cd in an environmental water sample. This results in electron flux through two electrochemical channels, displayed as ratio-metric heatmaps with real-time tracking over 42 hours (clean water) or 110 hours (environmental water sample). The red heatmap indicates the CymA-mediated channel, while the blue heatmap indicates the Flavin-mediated channel. The heatmaps are processed into 2-bit output signals based on a predetermined threshold (CymA > 1.15, Flavin > 1.2). The output signals accurately track the input signal (− for 0 and + for 1). Response times are determined when the 2-bit digit signal precisely represents the analyte conditions, shown in columns (dash filled for no analyte, light gray filled for one analyte, and dark gray for both analytes). Under clean water samples, inducer-E. coli could detect the presence of IPTG at 18 hours. The presence of aTc alone or both IPTG and aTc was detected at 30 hours. For heavy metal-E. coli, the presence of As or Cd alone was detected at 18 hours, while the presence of both heavy metals was detected at 30 hours. When under environmental water samples, the presence of both analytes or As alone was detected at 18 hours, while the presence of Cd alone was detected at 30 hours.

To test the applicability of our system in a real-world scenario, we evaluated our bioelectronic sensor in riverine samples with different combinations of As and Cd (Fig 4D, 4E). Environmental water samples were collected from Brays Bayou in Houston. We first removed the large solids by passing samples through a 0.2-micron filter paper, then added glycerol as a carbon source in M9 minimal medium (Fig 4D). We then exposed the engineered cells to the water samples with different combinations of heavy metals (2.5 μM As or/and 800 nM Cd). We continuously monitored the output signals using both single-potential and double-potential approaches for up to 110 hours. The current output at a single-potential reflects the electron transfer capacity of the heavy metal-*E. coli*. The current outputs obtained under single potential were much lower compared to those observed in clean water, indicating less electroactivity of engineered cells in the environmental water (Fig S7). However, the relative current ratios encoded by the double-potential process remain similar to those from clean water. The response time for all combinations of analytes in environmental water was by 27 hours or less, similar to the response time with clean water (Fig 4E). We can also detect certain combinations of analytes in a shorter response time. For instance, we can detect the presence of Cd or coexistence of As and Cd within 14 hours and identify the presence of As at the 27-hour mark in the environmental water sample. Our sensor continued to deliver a stable and reliable sensing output for up to 98 hours. However, beyond this timeframe, the signal output of (1,1) indicating the coexistence of As and Cd transitioned to (1,0), where the second digit to indicate Cd presence transitioned from ‘1’ to ‘0’ despite Cd remaining in solution. This inaccurate transition suggests a potential issue with prolonged analyte detection in this particular channel. In summary, our bioelectronic sensor provides stable and reliable continuous monitoring of heavy metals in complex environmental waters over a long-term period.

## DISCUSSION

Expanding the information capacity of individual bioelectronic sensors instead of relying on multiple sensors could greatly save on cost and space in advanced sensing applications. To achieve this, we successfully engineered two different EET pathways, the CymA and the Flavin synthesis pathway, into *E. coli* simultaneously. Each pathway is individually activated by promoters responsive to either the inducers IPTG or aTc, or the heavy metals As or Cd resulting in strains containing two analyte-sensitive electrochemical channels. Initially, there was difficulty in differentiating the current signals generated from the two EET pathways when recording at single potential. To solve this, we used double-step potentiometry and developed an algorithm capable of calculating the relative current ratios for each channel. When the ratios surpass the predefined threshold, the signal is converted from 0 to 1, indicating the presence of the desired analyte. In this way, a simple binary readout is generated from the two channels that records the presence of one analyte, the other analyte, or both analytes. Finally, the system was able to operate for days in both clean and environmental water samples. This platform facilitates precise and continuous monitoring of multiple analytes, enhancing the accuracy and effectiveness of bioelectronic sensors.

In addition to expanding the sensing capability to four states, our redox-potential-based algorithm works independent of specific signal receptors or EET pathways. The strains could be re-engineered with different promoters to sense other analytes, opening up a range of sensing targets, such as small-molecules^33^, physical changes^34^, and nucleic acids^35,36^. Moreover, the encoded signals involve both directed and mediated electron transfer processes, highlighting its versatility across specific electron transfer mechanisms. We envision that our algorithm also be employed for more than 2 channels, as long as the redox potentials of the EET pathways are distinguishable. This expands the range of various EET-associated molecules or proteins that could serve as channels, such as omcS^37^, phenazines^38^, and quinones^39^. Leveraging the well-established biosynthesis pathways of these redox proteins and mediators, we could integrate them with diverse signal receptors. By integrating variable sensing components and multiple channels with our agnostic algorithm, the capabilities of bioelectronic sensors could be significantly expanded, even reaching high-throughput levels.

However, the response time of our sensor is quite slow compared to other bioelectronic sensors. Our previous bioelectronic sensors were able to detect an endocrine disruptor within 2 minutes by incorporating a post-translationally controlled synthetic EET process. The much longer response time of 30 hours of our multichannel biosensor is mainly associated with the slow transcriptional regulation process. The longer response time could also be attributed to the time required to express a sufficient amount of protein to transfer enough electrons to be detectable. Further improvement could be achieved by integrating a post-translational regulation system into the sensor design.

Our study expands the boundaries of bioelectronic sensing. The two transcriptionally controlled electrochemical channels impart more information than previously designed bioelectronic sensors and the overall platform offers the flexibility to integrate diverse sensing systems and EET processes. This technology has applications for dynamic monitoring in complex conditions, such as polluted water environments and gastrointestinal tracts. By providing valuable insights into these complex systems, this technology can aid in effective environmental management strategies, disease diagnosis, and personalized healthcare interventions.

## CONCLUSIONS

By combining synthetic gene circuit design and electron transport in bacterial cells with electrochemical signal data processing, we developed a multichannel bioelectronic sensor capable of *in situ* sensing of multiple contaminants in environment samples over a timescale of days. Using heavy metal responsive promoters in urban water as a proof-of-concept model system, we demonstrate a flexible strategy for gene circuits design and signal optimization, and develop an agnostic encoding program for signal output.

## Supporting information

Support information

## ACKNOWLEDGEMENTS

The authors acknowledge the CAF Lab members for helpful discussion and Shyam Bhakta for input on the modular cloning approaches. We acknowledge the invaluable assistance of Marimikel Charrier, Biki Kundu and Matt Carpenter in proofreading the manuscript. Caroline Ajo-Franklin and Xu Zhang acknowledge supported by the Cancer Prevention and Research Institute of Texas (RR190063).

## METHODS AND MATERIALS

### Plasmids assembly and transformation

A list of all plasmids with primers and strains used in this study are provided in Supplementary Table S1. *E. coli* 5-a was used for all plasmid construction and amplification. The promoter sequences are provided table S2. The key primers and overhang sequences used for cloning are shown in table S2. The plasmids were constructed through Gibson assembly. All assembled plasmids were verified using sanger DNA sequencing. For experiments using *E. coli* cells, plasmids were transformed into chemically competent *E. coli* (DE3). *E. coli* was then plated on LB-agar plates containing a combination of 50 mg/L kanamycin and 34 μg/mL chloramphenicol and incubated overnight at 37.

#### Culturing conditions

We first inoculated all the *E. coli* strains directly from −80 °C glycerol stocks in 5 mL LB media with antibiotics (50 μg/mL kanamycin and 34 μg/mL chloramphenicol) and grew them at 37 °C overnight with 250 rpm sharking. Then we inoculated cell cultures (1:100 v/v ratio) in 50 mL 2xYT media in foil-covered 250 mL flasks with the same antibiotic concentrations and 1 mM δ-ALA (aminolevulinic acid) and grew these cultures at 37 °C with 250 rpm sharking.

#### Cells in vitro assay

When the cell OD600 reached between 0.5-0.6, we added IPTG, aTc, and heavy metals depending on plasmids used to induce the gene expression, followed by culturing at 30 °C with 250 rpm shaking overnight. Then we washed cells twice with M9 minimal medium.

#### Cells for bioreactors

The overnight culture also transferred into flasks (1:100 v/v ratio) with the same antibiotic concentration and δ-ALA. After the cell OD600 reached between 0.5-0.6, the cell was collected and washed with M9 minimal medium twice before inoculating the bioreactors.

#### MoO3 Nanoparticle assay to assess CymA pathway

Resuspended the washed cells to an OD600 of 1 with the M9 minimal medium including 1% glycerol. Resuspended cultures were mixed at a 1:1 ratio with 10 mg/mL molybdenum trioxide (MoO_3_) nanoparticles (Sigma-Aldrich) and incubated in a 96-well plate at 30 °Ce in a 5% H_2_/10% CO_2_/85% N_2_ atmosphere. When the blueness becomes stable, we record the cultures to represent the EET activities of CymA pathway, using a CanoScan LiDE 220 instrument, and the blueness of wells was determined using ImageJ with the ReadPlate3 plugin [ref]. MoO3 nanoparticles (<100 nm) were from Sigma.

#### Enhanced Chemiluminescence Staining (ECL)

Cells were pelleted (4000 × g) for 10 min at room temperature and washed with M9 minimal medium 3 times. Cultures having the same OD600 were mixed in a 1:1 ratio with a gel loading solution containing mercaptoethanol (0.71 M) and 1× LDS Loading Dye Mix (NuPage), and this mixture was incubated for 15 min at 95 °C. Samples were then loaded onto a 12% Bis-Tris gel (Invitrogen) and run for 45 min at 160 mV with a Precision Plus Protein Dual Color Standards ladder. Protein samples were transferred to blotting paper using a Trans-Blot Turbo Transfer Pack (Bio-Rad) and a Trans-Blot Turbo (Bio-Rad) using the “1 midi gel” setting. Cytochromes were visualized using Clarity Western ECL Substrate (Bio-Rad) and then imaged for both visible and chemiluminescent signals using a FluorChem E imager (ProteinSimple). Chemiluminescent images were overlaid on visible images to compare protein standards and chemiluminescent cytochromes.

#### Assessment of Flavins

Flavins have the intrinsic fluorescence properties related to concentration gradients. Fluorescence measurements were performed with 100 μL of experimental culture in a 96-well plate. The flavins were measured with 450/530 nm as excitation/emission wavelength. Optical density at 600 nm (OD600) was also measured as well.

#### Bioelectrochemical characterizations

We used two-chamber, 3-electrode electrochemical reactors to perform the electrochemical experiments as previously described^29^. The reactors contained three electrodes: a working electrode (graphite felt, diameter of 3.2 cm, surface area 16 cm2 for two sides, Alfa Aesar) and a reference electrode (CHI111, Ag/AgCl with 3 M KCl, CH Instruments) in the anodic chamber, and a counter electrode (titanium wire, Alfa Aesar) in the cathodic chamber. We also used a cation exchange membrane (CMI-7000S, Membranes International) to separate these two chambers. Each chamber contained 120 mL M9 minimal medium (BD) and 1% glycerol (Sigma-Aldrich) as the electrolyte. The anodic chamber was continuously stirred with a stir bar at 300 rpm, temperature-controlled at 30 °C. BioLogic-potentiostat were used to conduct all the electrochemical measurements (VSP-300, Bio-Logic Science Instruments). Single-potential chronoamperometry was conducted by applying potential at 0.2 V vs. Ag/AgCl with the current recorded every 30 s. To differentiate two electrochemical channels, we applied a double potential step with current recording over a 30 min decay for each potential step. For the CymA channel, we polarized the electrode at 0.2 V and 0 V vs Ag/AgCl, while for the Flavin channel the electrode potentials were at -0.2 V and -0.4 V vs. Ag/AgCl.

## AUTHOR CONTRIBUTIONS

All authors contributed to the design of experiments, collecting of data, and writing of the manuscript.

