## Supplementary material for "Multichannel bioelectronic sensing using engineered *Escherichia coli*": Support information

Support information for:

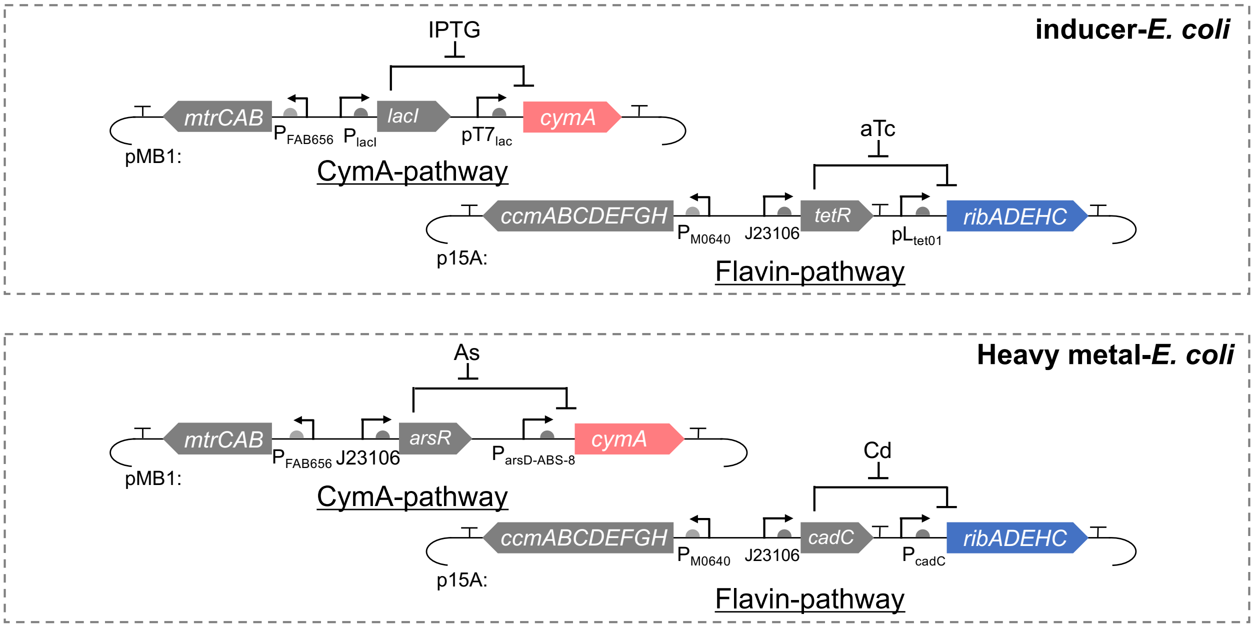


**Fig S1.** Schematic of the genetic circuits for EET pathways with different sensing systems.

**Supplementary Table 1. All plasmids used in this study.**

KanR: kanamycin resistance gene, CmR: chloramphenicol resistance gene, pMB01: origin of replication, p15A: origin of replication, pT7: T7 promoter. CymA: gene of interest, MtrCAB: extracellular electron transfer pathway, ccmABCDEFGH: cytochrome c maturation (Ccm) system, ribADEHC: flavin synthesis pathway.

| Name | Description | Plasmid details |
|---|---|---|
| **pMB01** | CymA expressing under IPTG regulation | lacI-pT7-CymA-pFAB656-MtrCAB-KanR-pMB01 |
| **p15A01** | Flavin synthesis expressing plasmid under aTc regulation | tetR-pLtet01-ribADEHC-pM0640-ccmABCDEFGH-CmR-p15A |
| **pMB07** | CymA expressing under As regulation | ArsR-ParsD-ABS-8-CymA-pFAB656-MtrCAB- KanR-pMB01 |
| **P15A02** | Flavin synthesis expressing plasmid under Cd regulation | CadC-pcadC-ribADEHC-pM0640-ccmABCDEFGH-CmR-p15A |

**Supplementary Table 2. DNA sequence for heavy metal responsive promoters**

| Name | DNA sequence |
|---|---|
| **arsR gene from *E.coli*** | ATGTCATTTCTGTTACCCATCCAATTGTTCAAAATTCTTGCTGATG<br>AAACCCGTCTGGGCATCGTTTTACTGCTCAGCGAACTGGGAGAG<br>TTATGCGTCTGCGATCTCTGCACTGCTCTCGACCAGTCGCAGCC<br>CAAGATCTCCCGCCACCTGGCATTGCTGCGTGAAAGCGGGCTAT<br>TGCTGGACCGCAAGCAAGGTAAGTGGGTTCATTACCGCTTATCA<br>CCGCATATTCCAGCATGGGCGGCGAAAATTATTGATGAGGCCTG<br>GCGATGTGAACAGGAAAAGGTTCAGGCGATTGTCCGCAACCTGG<br>CTCGACAAAACTGTTCCGGGGACAGTAAGAACATTTGCAGTTAA |
| **ParsD-ABS-8** | TGTTTTTGACTTATCCGCTTCGAAGAGAGATACTTACACATTCGTT<br>AAGTCATATATGTTTTTGACTTATCCGC |
| **CadC gene from P.putida** | ATGAAGAAGAAGGATACCTGCGAGATCTTCTGCTACGACGAGGA<br>AAAGGTGAACCGCATCCAGGGCGACCTGCAGACCGTGGACATC<br>AGCGGTGTGAGCCAGATCCTGAAGGCGATCGCGGACGAGAACC<br>GTGCCAAGATCACCTACGCGCTGTGCCAGGACGAGGAACTGTG<br>CGTGTGCGATATCGCCAACATCCTGGGCGTGACCATCGCCAACG<br>CCAGCCACCATCTGCGCACCCTGTACAAGCAGGGCGTCGTGAA<br>CTTCCGCAAAGAGGGCAAGCTGGCCCTGTACAGCCTGGGCGAC<br>GAGCACATCCGCCAGATCATGATGATCGCCCTGGCGCACAAGAA<br>AGAGGTGAAGGTCAACGTCTGA |
| **pCadC** | CTTTTATTTTCATTCAAATATTTGCTTGCATGATGAGTCGAAAATG<br>GTTATAATACACTCAAATAAATATTTGAAT |

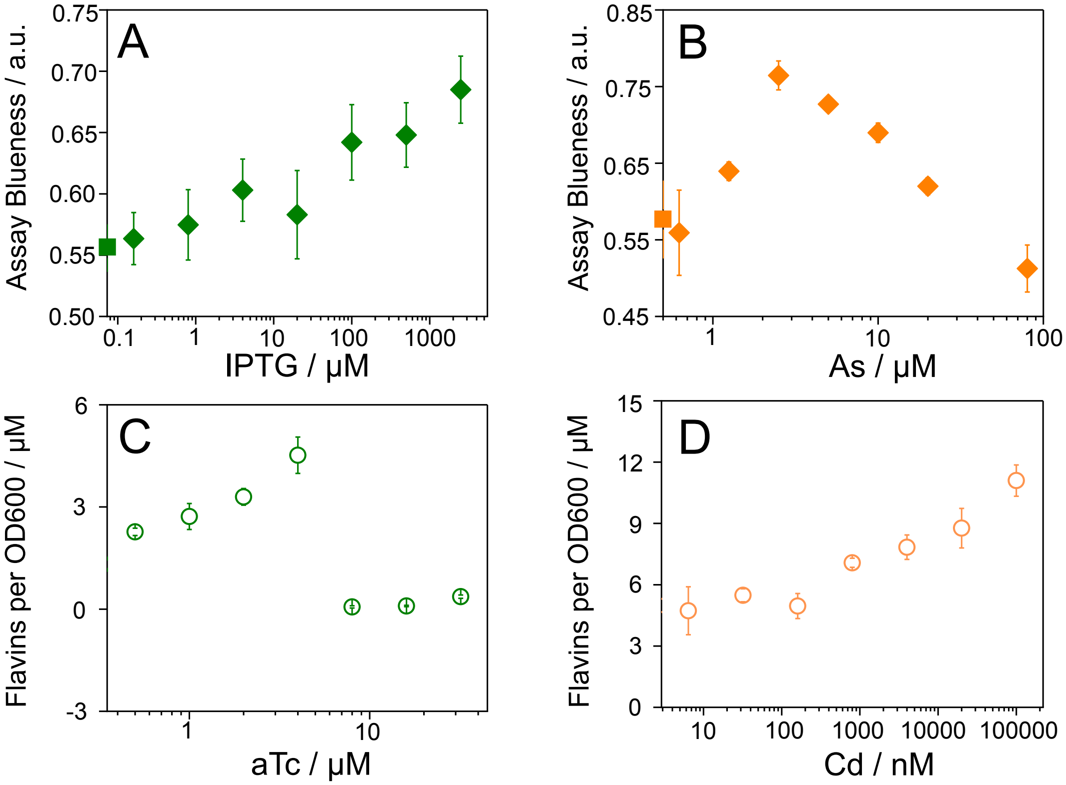


**Fig S2.** Characterization of individual EET pathways under different analyte concentration. A or B, in vitro evaluation of the expression level of CymA under IPTG or As regulation, using a colorimetric molybdenum-based nanoparticle assay; C or D, extracellular flavin production as the function of aTc or Cd concentration, representing the flavin synthesis pathways expression levels. Scatters show the mean values and error bars represent standard deviation of n=4 biological replicates.

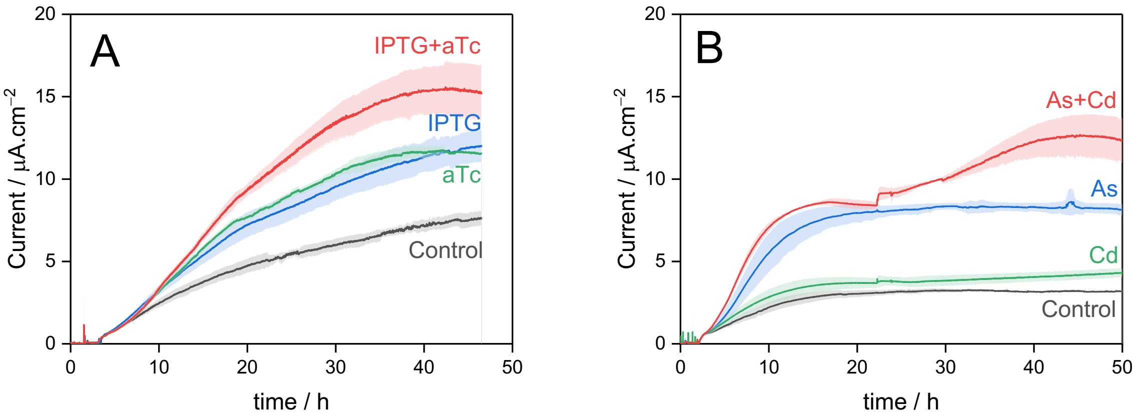


**Figure S3.** The electrical current signals at a constant potential at 0.2 V vs Ag/AgCl from two engineered strains A) inducer-E. coli and B) heavy metal-E. coli. The different colors indicate different analytes conditions.

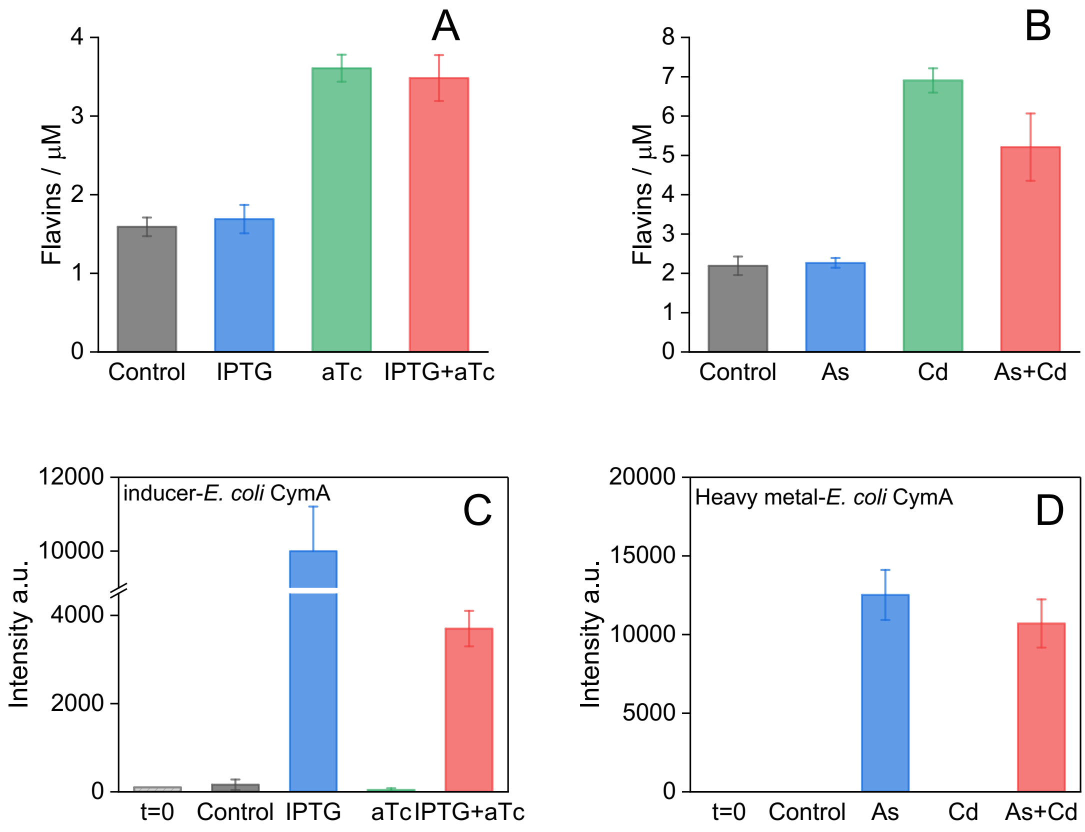


**Figure S4.** The evaluation of extracellular flavin concentrations and CymA expression in response to analytes for inducer-*E. coli* and heavy metal-*E. coli*. Samples collected at the end of the operational process, columns show the mean values, and error bars represent standard deviation of n=3 biological replicates.

**Double Step Chronoamperometry:**

Taking CymA pathway as example where its midpoint potential is around 0 mV vs Ag/AgCl, where we have applied double potentials where localized 200mV as oxidation dominance status, and 0 mV as the equilibrium. Assume that the mass transfer is diffusion driven process, we can simplify the Nernst-Plank equation as:

$$J_i(x) = -D_i \frac{\partial C_i(x)}{\partial x}$$

Where $J_i(x)$ is the flux of species I (mol.$s^{-1}cm^{-2}$) at distance x from the surface, Di is the diffusion coefficient ($cm^2/s$), $\partial C_i(x)/\partial x$ is the concentration gradient at distance x.

Assuming the rate of mass transfer is proportional to the concentration of gradient at the electrode surface, as given by the diffusive part

$$v_{mt} \propto (dC_O/dx)_{x=0} = D_O(dC_O/dx)_{x=0}$$

Assume a linear concentration gradient within diffusion layer, then from equation,

$$v_{mt} = D_O[C_O^* - C_O(x = 0)]/\delta_O$$

Since diffusion layer is often unknown, it is convenient to combine it with the diffusion coefficient to produce a single constant, $m_O = D_O/\ \delta_O$

$$v_{mt} = m_O(C_O^* - C(x = 0)$$

The proportionality constant, $m_O$, called the mass-transfer coefficient, has units of cm.$s^{-1}$. Since $C_O^* > C_O(x = 0)$

$$\frac{i}{nFA} = m_O[C_O^* - C_O(x = 0)]$$

The value of $C_O(x = 0)$ are function of electrode potential E. The largest rate of mass transfer of CymA occurs when $C_O(x = 0) \ll C_O^*$. The value of current under these conditions is called the limiting current, $i_l$, where

$$i_l = nFAm_OC_O^*$$

Which indicates that the current values obtained at steady-state current is proportional to the concentration of redox species. When IPTG or As is present in the solution, resulting in higher concertation of CymA expression, which increase $C_O^*$, the ratio of two steady-state currents can indicate the concentration levels of CymA expression.

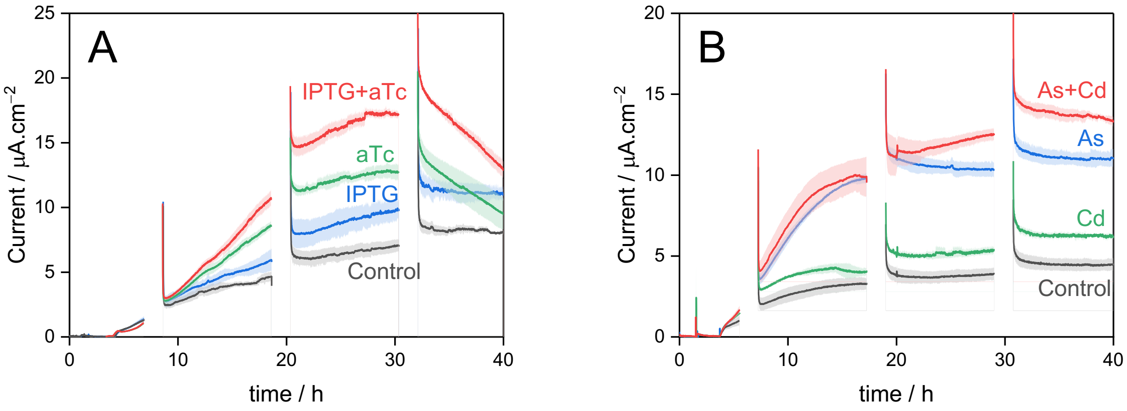


**Figure S5.** The electrical current signals at a constant potential of 0.2 V vs. Ag/AgCl from two engineered *E. coli* strains A) inducer-*E. coli* and B) heavy metal-*E. coli*. Intermittent gaps highlight the sampling intervals where double potential step chronoamperometry was employed, and by using redox-potential-dependent algorithms, to differentiate sensing signals in response to various analytes. Lines show the mean values and error bars represent standard deviation of n=3 biological replicates.

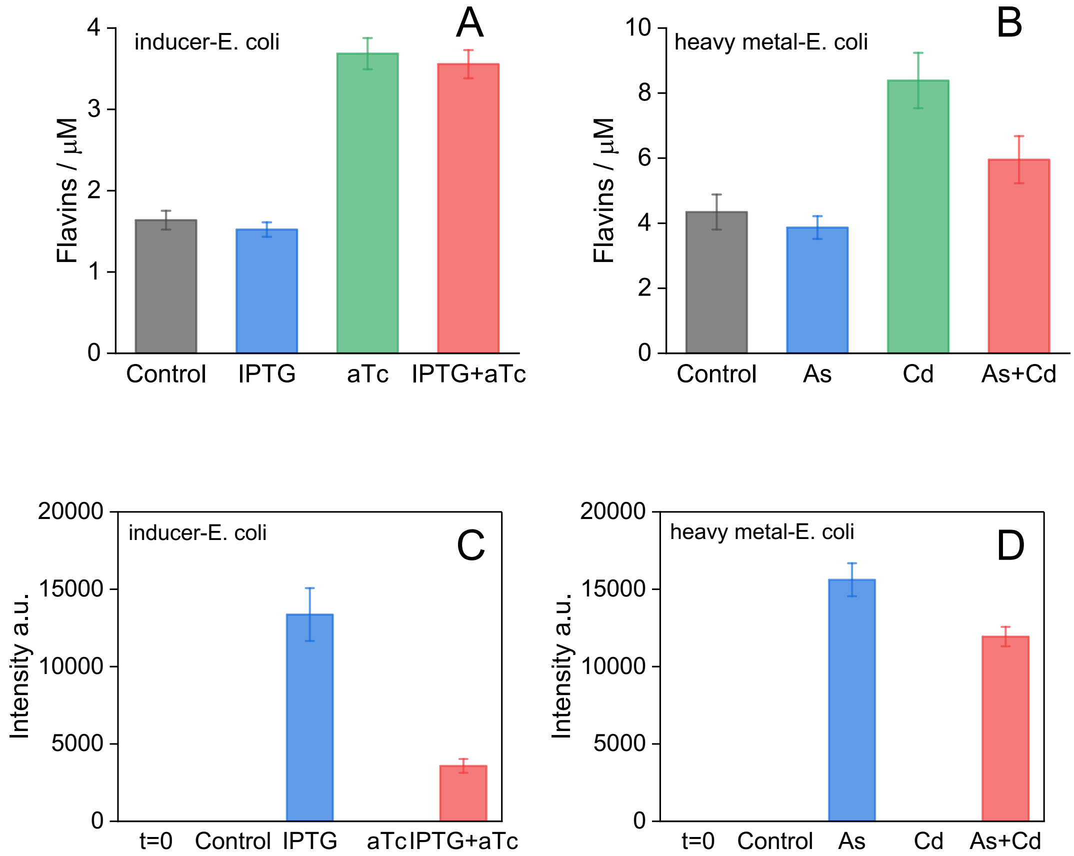


**Figure S6.** The evaluation of extracellular flavin concentrations and CymA expression in response to various analytes after continuous monitoring of algorithmically processed signals. Extracellular flavin concentration under different conditions for A) inducer-*E. coli* strain and B) heavy metal-*E. coli* strain. Expression level of CymA under different conditions for C) inducer-*E. coli* strain and D) heavy metal-*E. coli* strain. Samples collected at the end of the operational process, columns show the mean values, and error bars represent standard deviation of n=3 biological replicates. The columns labeled as t=0 represent the cell samples of the initial inoculation.

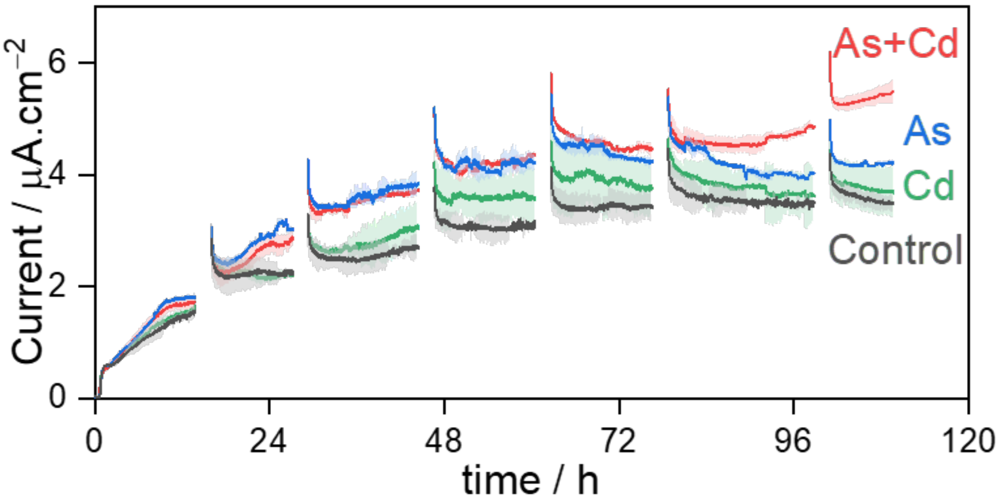


**Figure S7.** The electrical current signals at constant potential of 0.2 V vs. Ag/AgCl from heavy metal-*E. coli* exposed to different analytes conditions in environmental water samples. Intermittent gaps highlighting the sampling intervals where double potential step chronoamperometry was employed, and by using redox-potential-dependent algorithms, to differentiate sensing signals in response to various analytes. Lines show the mean values and error bars represent standard deviation of n=3 biological replicates.

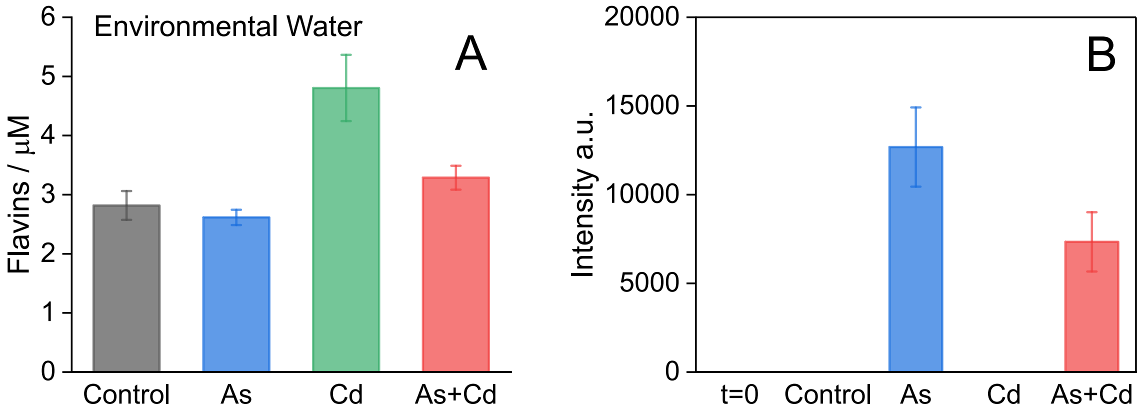


**Figure S8.** The evaluation of extracellular flavin concentrations and CymA expression of heavy metal-*E. coli* in response to various analytes with environmental water samples. Samples collected at the end of the operational process, columns show the mean values, and error bars represent standard deviation of n=3 biological replicates. The columns labeled as t=0 represent the cell samples of the initial inoculation.
